# A protocol for single-cell transcriptomics from cryopreserved renal tissue and urine for the Accelerating Medicine Partnership (AMP) RA/SLE network

**DOI:** 10.1101/275859

**Authors:** Deepak A. Rao, Celine C. Berthier, Arnon Arazi, Anne Davidson, Yanyan Liu, Edward P. Browne, Thomas M. Eisenhaure, Adam Chicoine, David J. Lieb, Dawn E. Smilek, Patti Tosta, James A. Lederer, Michael B. Brenner, David Hildeman, E. Steve Woodle, David Wofsy, Jennifer H. Anolik, Matthias Kretzler, Nir Hacohen, Betty Diamond

**Affiliations:** Division of Rheumatology, Allergy, Immunology, Department of Medicine, Brigham and Women’s Hospital, Harvard Medical School, Boston MA 02115; Internal Medicine, Department of Nephrology, University of Michigan, Ann Arbor, MI, USA 48109; Broad Institute of MIT and Harvard, Cambridge MA 02142; Center for Autoimmune and Musculoskeletal Diseases, The Feinstein Institute for Medical Research, Northwell Health, Manhasset, NY 11030; Lupus Nephritis Trials Network, University of California San Francisco, San Francisco, CA 94107; Immune Tolerance Network, University of California San Francisco, San Francisco, CA 94107; Rheumatology Division and Russell/Engleman Research Center, University of California San Francisco, San Francisco, CA 94107; Department of Surgery, Brigham and Women’s Hospital, Harvard Medical School, Boston MA 02115; Department of Pediatrics, University of Cincinnati, Cincinnati, OH 45257; Division of Immunobiology, Cincinnati Children’s Hospital Medical Center, Cincinnati, OH 45229; Section of Transplantation, Department of Surgery, University of Cincinnati College of Medicine, Cincinnati, OH 45257; University of Rochester Medical Center, Rochester, NY 14642; Center for Cancer Research, Massachusetts General Hospital, Harvard Medical School, Boston MA 02114

## Abstract

**OBJECTIVE:** There is a critical need to define the cells that mediate tissue damage in lupus nephritis. Here we aimed to establish a protocol to preserve lupus nephritis kidney biopsies and urine cell samples obtained at multiple clinical sites for subsequent isolation and transcriptomic analysis of single cells.

**METHODS:** Fresh and cryopreserved kidney tissue from tumor nephrectomies and lupus nephritis kidney biopsies were disaggregated by enzymatic digestion. Cell yields and cell composition were assessed by flow cytometry. Transcriptomes of leukocytes and epithelial cells were evaluated by low-input and single cell RNA-seq.

**RESULTS:** Cryopreserved kidney tissue from tumor nephrectomies and lupus nephritis biopsies can be thawed and dissociated to yield intact, viable leukocytes and epithelial cells. Cryopreservation of intact kidney tissue provides higher epithelial cell yields compared to cryopreservation of single cell suspensions from dissociated kidneys. Cell yields and flow cytometric cell phenotypes are comparable between cryopreserved kidney samples and paired kidney samples shipped overnight on wet ice. High-quality single cell and low-input transcriptomic data were generated from leukocytes from both cryopreserved lupus nephritis kidney biopsies and urine, as well as from a subset of kidney epithelial cells.

**CONCLUSION:** The AMP RA/SLE cryopreserved tissue analysis pipeline provides a method for centralized processing of lupus nephritis kidney biopsies and urine samples to generate robust transcriptomic analyses in multi-center studies.

---

Lupus nephritis (LN) is a frequent complication of systemic lupus erythematosus (SLE) that confers substantial morbidity and mortality(1-3). Current therapies are both toxic and insufficiently effective, with a substantial number of patients progressing to end stage renal disease and death(4,5). Despite the rapid pace of immunologic discovery, most clinical trials of rationally designed therapies have failed in both general SLE and LN, and only one new drug has been approved for the treatment of lupus in the last 5 decades(6). To alter the course of this disease, there is a pressing need to refocus our efforts on defining mechanisms that drive target organ damage and are amenable to intervention with novel or repurposed therapeutic agents.

In recognition of this need, the National Institutes of Health (NIH), the pharmaceutical industry, and non-profit organizations joined together in 2014 to form the Accelerated Medicine Partnership (AMP) RA/SLE network(7), whose goal is to identify new diagnostic and therapeutic targets through a better understanding of the mechanisms by which individual cell types contribute to tissue damage. The AMP has deployed a staged approach with technology development in Phase 0, initial field testing in a pilot LN cohort in Phase 1, and a large scale clinical study using the optimized protocol in Phase 2.

One of the central goals of the AMP RA/SLE network is to acquire and analyze kidney biopsy samples from patients with LN by single cell transcriptomics. Such analyses performed on a large population of patients have the potential to identify critical inflammatory pathways that define subsets of patients with distinct pathophysiologic features. These data may allow for the development of predictive biomarkers for personalized therapeutic approaches and for discovery of novel therapeutic targets. Acquisition of a sufficient number of samples and generation of robust, reproducible data within a reasonable time frame represent central challenges for the AMP consortium. To this end, we aimed to develop a standardized protocol for acquisition of the LN samples across a distributed research network. A further goal was to enable central processing and analysis at the AMP technology sites to minimize inter-site variability and batch effects.

During the AMP Phase 0, we considered 3 options for centralized analysis of samples acquired at multiple sites: 1) dissociation of tissue into single cells at the acquisition site, followed by cryopreservation of the single cell suspension, 2) overnight shipment of tissue samples in a storage solution, or 3) immediate cryopreservation of the tissue sample, followed by dissociation at a central processing site. Requiring individual acquisition sites to dissociate samples in order to cryopreserve single cell suspensions limits the number of collection sites to only those capable of sufficiently sophisticated laboratory bench work, and may introduce inter-site variability in tissue processing technique. Cryopreservation may also introduce selective cell losses. Alternatively, analysis of samples shipped in a storage solution on wet ice may introduce artefacts of overnight shipping. In addition, overnight shipping imposes a considerable logistical challenge; because the tissue is not stable, the centralized processing site must be available to process samples immediately upon arrival. The strategy of immediate cryopreservation of intact kidney tissue followed by batch transportation offered a simple, accessible workflow for acquisition sites, as well as flexible sample processing in batches at a centralized site to minimize technical variation. We therefore evaluated the feasibility and performance of this strategy, compared to the alternative approaches.

Here, we describe a protocol to dissociate and analyze viable immune and epithelial cells from cryopreserved kidney biopsies by flow cytometric cell sorting and low-input and single cell transcriptomics. We report that tissue leukocytes extracted from cryopreserved kidney tissues remain largely viable and yield robust global transcriptomes. A subset of epithelial cells obtained from frozen tissue is also of adequate quality for transcriptomic analyses. In addition, we find that cryopreserved cells from urine provide robust transcriptomic data, enabling paired analyses of leukocyte populations in kidney and urine. These methods are adaptable for use in multiple sites and have been adopted and utilized by the AMP RA/SLE network.

## MATERIALS AND METHODS

### Human kidney tissue and urine acquisition

Renal tissue and urine specimens were acquired at 4 clinical sites in the United States. The study received institutional review board approval at each site. Two types of human kidney tissue were collected. For initial development of the kidney processing method, kidney tissue samples were obtained from a tumor-free part of nephrectomy specimens (TN) and were transported to the laboratory on ice in HypoThermosol FRS Preservation Solution (BioLife Solutions).

For validation of the methods developed with tumor nephrectomies, research biopsy cores were acquired during the course of diagnostic renal biopsies performed on participants with LN as part of routine clinical care and were stored in HypoThermosol for transport as above. Midstream urine samples were collected and transported to the laboratory in sterile containers.

### Kidney cryopreservation

Each TN specimen was placed in a 10 ml centrifuge tube containing 5 ml of cold HypoThermosol FRS preservation solution and placed on ice. TN samples were then cut into thin segments with mass typically under 6 mg to mimic the size of a kidney biopsy specimen. Each LN biopsy was placed in a 1.5 ml centrifuge tube containing 1 ml of cold HypoThermosol FRS preservation solution. The tube was placed on ice until the sample was processed (typically ~20 minutes after biopsy). For cryopreservation, each piece was weighed and transferred into a cryovial containing 1 ml of CryoStor CS10 Freeze Media (BioLife Solutions). The cryovial was incubated on ice for 20-30 min to allow the solution to penetrate the tissue, and was then placed in a cold Mr. Frosty freezing container (Nalgene, #5100-0001) at -80°C overnight. Cryopreserved samples were then stored for up to 60 days in liquid nitrogen and then shipped on dry ice to the central processing site, where they were stored in liquid nitrogen until use.

### Kidney tissue thawing and dissociation into a single cell suspension

The cryovial containing the kidney tissue (TN or LN biopsy) was rapidly warmed in a 37°C water bath until almost fully thawed. The sample was then poured into a well of a 12- or 24-well dish and rinsed in a second well containing warmed RPMI/10%FBS. The biopsy was incubated for 10 minutes at room temperature.

To increase surface area, specimens were cut into 2-3 pieces and placed into a 1.5 ml centrifuge tube containing 445uL of Advanced DMEM/F-12 (ThermoFisher Scientific, #12634-028) and 5uL of DNase I (Roche, #04536282001, 100U/ml final concentration). 50uL of Liberase TL (Roche, #05401020001, 250ug/mL final concentration) was added and the tube was placed on an orbital shaker (300-500 rpm) at 37°C for 12 minutes. At 6 minutes into the digestion, the mixture was gently pipetted up and down several times using a cut 1 mL pipette tip. After 12 minutes, 500uL of RPMI/10% FBS was added to stop the digestion. Each resulting cell suspension was filtered through a 70-μm filter into a new 1.7 ml microfuge tube. An additional 200μL of RPMI/10%FBS was used to wash out the 1.7 mL microfuge tube in which the digestion took place. The remaining undigested tissue was pushed through the 70-um filter using the plastic end of a plunger of a 3 mL syringe, and an additional 200μL of RPMI/10%FBS was used to rinse the filter. The cell suspension was centrifuged at 300g at 4°C for 10 min. The supernatant was aspirated off, and the cell pellet was resuspended in 50-80uL of RPMI/10%FBS. Cells were then transferred to a 5 mL FACS tube and resuspended in cold PBS for downstream analyses.

### Urine cell pellet collection and cryopreservation protocol

The total urine volume (15-90 mL) was split into two 50 mL Falcon tubes. Urine cells were pelleted by centrifugation at 200g for 10 minutes, and then resuspended in 1 ml of cold X-VIVO10 medium (Lonza BE04-743Q). Cells were transferred to a microcentrifuge tube, washed once in 1mL of X-VIVO10 medium, and then resuspended in 0.5 mL of cold CryoStor CS10. Cells were transferred into a 1.8 mL cryovial, placed in a Mr. Frosty freezing container, stored in at -80°C overnight, and then transferred to liquid nitrogen. For downstream analyses, cryopreserved urine cells were rapidly thawed by vigorous shaking in a 37°C water bath, transferred into warm RPMI/10%FBS, centrifuged at 300g for 10 minutes, and resuspended in cold HBSS/1%BSA.

### Flow cytometric cell sorting

An 11-color flow cytometry panel was developed to identify epithelial cells and leukocyte populations within dissociated kidney cells. Antibodies include anti-CD45-FITC, anti-CD19-PE, anti-CD11c-PerCP/Cy5.5, anti-CD10-BV421, anti-CD14-BV510, anti-CD3-BV605, anti-CD4-BV650, anti-CD8-BV711, anti-CD31-AlexaFluor700, anti-PD-1-APC, and propidium iodide (all from BioLegend). Kidney or urine cells were incubated with antibodies in HBSS/1%BSA for 30 minutes. Cells were washed once in HBSS/1%BSA, centrifuged, and passed through a 70 micron filter.

Cells were sorted on a 3-laser BD FACSAria Fusion cell sorter. Intact cells were gated according to FSC-A and SSC-A. Doublets were excluded by serial FSC-H/FSC-W and SSC-H/SSC-W gates. Non-viable cells were excluded based on propidium iodide uptake. Cells were sorted through a 100 micron nozzle at 20 psi. For low-input bulk sorting, up to 4 populations were simultaneously sorted directly into 350uL of RLT lysis buffer (Qiagen) containing 1% β-mercaptoethanol. For single cell RNA-seq, cells were sorted into 96-well plates containing 5μL of TCL buffer (Qiagen) containing 1% β-mercaptoethanol. For comparisions to intact tissue, tissue samples were placed in 350μL RLT lysis buffer containing 1% β-mercaptoethanol and homogenized using a TissueLyser II (Qiagen).

### Quantification of cell yield

Quantification of cell yields from these small tissue samples was performed by hemocytometer with trypan blue exclusion and by flow cytometry with propidium iodide-exclusion. Yields of cell subsets (leukocytes, epithelial cells) were quantified by acquiring the entire sample through the flow sorter and plotting the number of intact, PI-negative cell events with the appropriate surface markers. Cell yields were normalized to input tissue mass.

### RNA extraction, library preparation and RNA-seq

RNA from sorted bulk cell populations was isolated using RNeasy columns (Qiagen). RNA from up to 1000 cells was treated with DNase I (New England Biolabs), then concentrated using Agencourt RNAClean XP beads (Beckan Coulter). Full-length cDNA and sequencing libraries were prepared using the SmartSeq2 protocol (Illumina) as previously described(8). Libraries were sequenced on a MiSeq (Illumina) to generate 25 base pair, paired-end reads. Single cell RNA-seq was performed using a modified SmartSeq2 protocol as described(9). Libraries were sequenced on a Hiseq 2500 (Illumina) in Rapid Run Mode to generate 76 base pair, paired-end reads.

### RNA-seq data analysis

Mapping of reads to human reference transcriptome hg19 and quantification of mRNA expression levels was done using RSEM(10). The percentage of reads mapped to transcripts and the number of genes detected were calculated. Identification of differentially expressed genes was performed using the Bioconductor package EdgeR(11), employing generalized linear models, in order to take into account the contributions of different variables such as the digestion enzyme used and freezing medium. The resulting p-values were adjusted using Benjamini and Hochberg’s procedure for controlling the false discovery rate (FDR)(12). Enriched GO terms in the lists of differentially expressed genes were found using Gorilla(13). Basic analyses of single cell RNA-seq data, including quantification of genes detected per cell, filtering of cells with less than 1,000 detected genes, and principal components analysis, were performed using Seurat(14).

### Assessing the quality of RNA-seq Libraries

Two main quality metrics were taken into account: the percentage of reads mapped to transcript, and the number of genes detected in the data by being associated with at least one read; the latter metric represents the complexity of the RNA-seq library. To allow comparison across samples, we normalized the number of detected genes by setting the total number of reads allocated to each sample to a fixed value (100K reads). To do this analytically, we took advantage of the next observation: if a gene has *n* reads mapped to it in the original data set, and a fraction *r* (0 ≤ *r* ≤ 1) of these reads is randomly chosen in order to get 100K reads, then the probability of having at least one of the *n* reads in the downsampled data is equal to 1 – (1 – *r*)^n^. We then summed up these probabilities in order to determine the average number of genes detected in the downsampled data.

### Classification of Sorted Urine Samples

To verify that the RNA-seq data obtained from sorted populations of urine cells contained a meaningful signal, the similarity of the sorted cell populations (CD45^+^CD3^+^,CD45^+^CD3^-^, CD45^-^) to a set of reference samples of known cell types was evaluated. For reference samples, we used a gene by sample expression matrix derived from FANTOM5 by computing the median expression level per gene, taken across the replicates corresponding to each of the 360 cell types included in this dataset. The similarity of each urine sample to each FANTOM5 samples was assessed by calculating the Pearson correlation between them. In order to take into account genes expressed at low levels while minimizing the noise associated with such measurements, we set each expression value lower than the first quantile (computed over positive values only) to the first quantile of the distribution.

### Statistical analysis

For protocol development, limited samples sizes per study were used in order to efficiently identify major effects that would substantially influence the final protocol. Individual data points are shown for each experiment. Statistical tests were applied for experiments with samples sizes greater than 4 using non-parametric tests, except for RNA-seq differential expression analyses described above.

## RESULTS

The goal of Phase 0 pipeline development was to generate a protocol for robust transcriptomic data generation from viable cells derived from LN kidney and urine samples. During development and optimization of this protocol, alternative approaches were tested for several of the key tissue processing steps. In the following sections, we report the main parameters considered.

### Effect of cryopreservation on cells dissociated from kidney tissue

Use of cryopreserved kidney cell samples offers the potential to analyze kidney biopsies acquired at multiple sites in a uniform, centralized manner. To evaluate the effect of cryopreservation on kidney cells, we first quantified cell yields from enzymatically-dissociated tumor nephrectomy samples before and after cryopreservation. Cryopreservation of dissociated kidney cells led to a significant reduction in cell yield (**Fig. 1a**, n=76 fresh samples, n=24 frozen samples).

**Figure 1.**
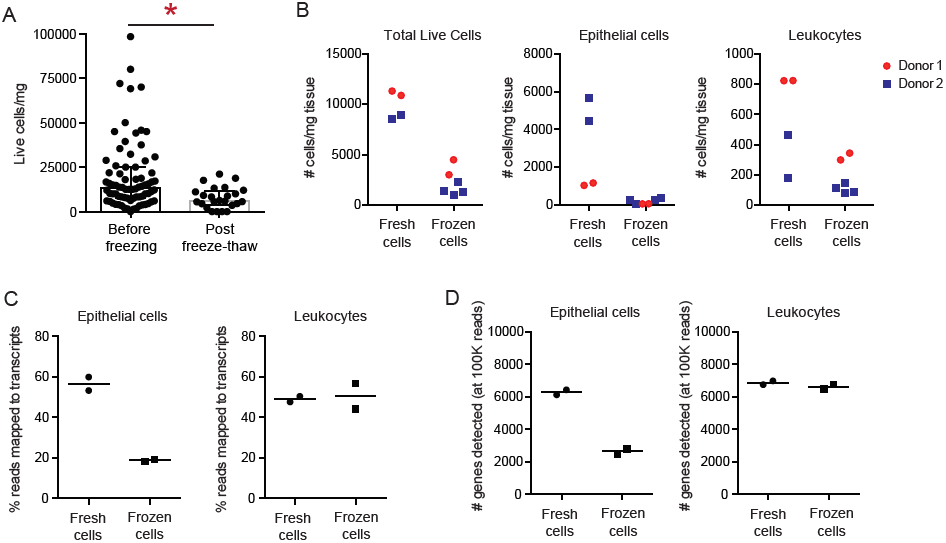
Cryopreservation of dissociated kidney single cells reduces viability, yield and transcriptomic data quality of epithelial cells, but not leukocytes. A) Total live cell yields by hemocytometer from kidney samples before or after freeze/thaw of dissociated cells. Before freeze n=76, post freeze/thaw n=24, * p=0.0009 by Mann-Whitney. B) Cell yields quantified by flow cytometry from tumor nephrectomy kidney tissue analyzed immediately after dissociation (fresh cells) or after freeze/thaw of dissociated cells (frozen cells). C,D) Percent of reads mapped to transcripts (C) or number of genes detected (D) in RNA sequencing data from tumor nephrectomy kidney cells sorted immediately or after freeze/thaw of dissociated cells (n=2).

Flow cytometric analyses revealed that epithelial cells were substantially diminished upon thawing, while leukocyte populations remained relatively preserved (**Fig. 1b**, n=2 donors with 2-4 replicates). Consistent with these findings, RNA-seq of libraries prepared from cryopreserved dissociated kidney cells demonstrated a considerable reduction in quality due to cryopreservation of epithelial cells, while transcriptomes of leukocytes sorted after cryopreservation were comparable to those of freshly isolated leukocytes (**Fig. 1c, d**, n=2). Consistent with prior reports, a transcriptomic signature suggesting cell stress could be detected in cryopreserved leukocytes, with upregulation of genes associated with such GO terms as ‘response to unfolded protein’, ‘response to stress’ and ‘response to temperature stimulus’, including heat shock proteins (Table S1) (15-17). Similar results were seen in kidney leukocytes from TN samples dissociated and frozen in two different cryopreservation media, CryoStor10 and human serum supplemented with 10% DMSO (Supplemental Figure 1).

### Cryopreservation of intact kidney tissue vs. dissociated cells

Because cryopreservation of intact tissue, rather than dissociated cells, offers a far simpler workflow for tissue acquisition sites, we compared cell yields from these two processing strategies. Flow cytometric quantification revealed that cryopreservation of intact TN tissue in CryoStor10 freezing solution consistently yielded more epithelial cells compared to cryopreservation of dissociated cells (**Fig. 2a**, n=9 segments from 4 different donors). Leukocyte yields were comparable between the two methods, with slightly higher yields from cryopreserved intact tissue. To assess whether cryopreserved intact tissue can yield cells adequate for downstream transcriptomic analyses, we sequenced cDNA libraries from samples prepared using either method. The quality of the libraries was found to be similar for leukocytes, while a somewhat higher quality was observed for epithelial cells when intact tissue, rather than a dissociated cell suspension, was cryopreserved (**Fig. 2b, c**, n=2).

**Figure 2.**
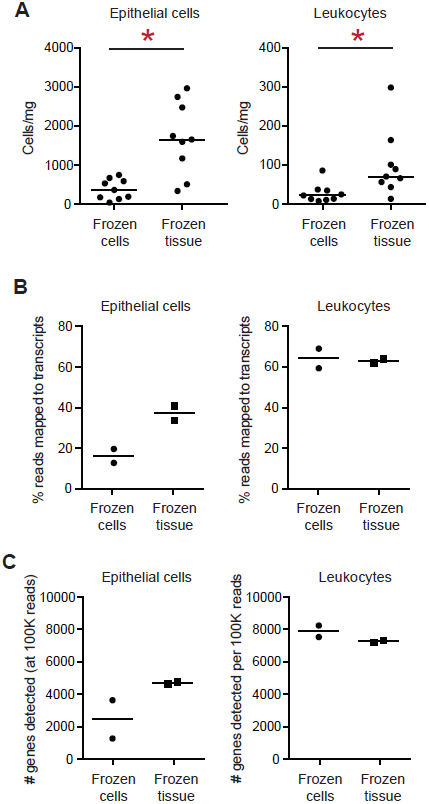
Cryopreservation of intact kidney tissue, rather than dissociated cells, improves epithelial cell recovery. A) Cell yields determined by flow cytometry from tumor nephrectomy kidney tissue either cryopre-served post-dissociation (frozen cells) or pre-dissociation (frozen tissue). n=9 segments pooled from 4 different donors (1-4 segments per donor). * p<0.01 by Mann-Whitney test. B,C) Percent of RNAseq reads mapped to transcripts (B) and number of genes detected (C) in RNA sequencing data from tumor nephrectomy kidney samples cryopreserved post-dissociation (frozen cells) or pre-dissociation (frozen tissue) (n=2 donors in separate experiments).

### Comparison of cryopreserved kidney tissue with kidney tissue shipped overnight on ice

We next compared cell yields from paired samples from 5 different donors analyzed either as cryopreserved whole tissue or as tissue shipped overnight in the HypoThermosol preservation solution on wet ice. Epithelial and leukocyte yields quantified by flow cytometry were variable but not consistently higher by either method (**Fig. 3a**, n=5). Flow cytometric detection of total CD45^+^ leukocytes, CD3^+^ T cells, and CD14^+^ monocyte/macrophages appeared generally similar with both methods, suggesting that cryopreservation does not substantially alter the frequency of these cell populations (**Fig. 3b**, n=5). The quality of RNA-seq libraries prepared from both leukocytes and epithelial cells from cryopreserved intact tissue was comparable to that from overnight-shipped samples (**Fig. 3c, d**, n=2).

**Figure 3.**
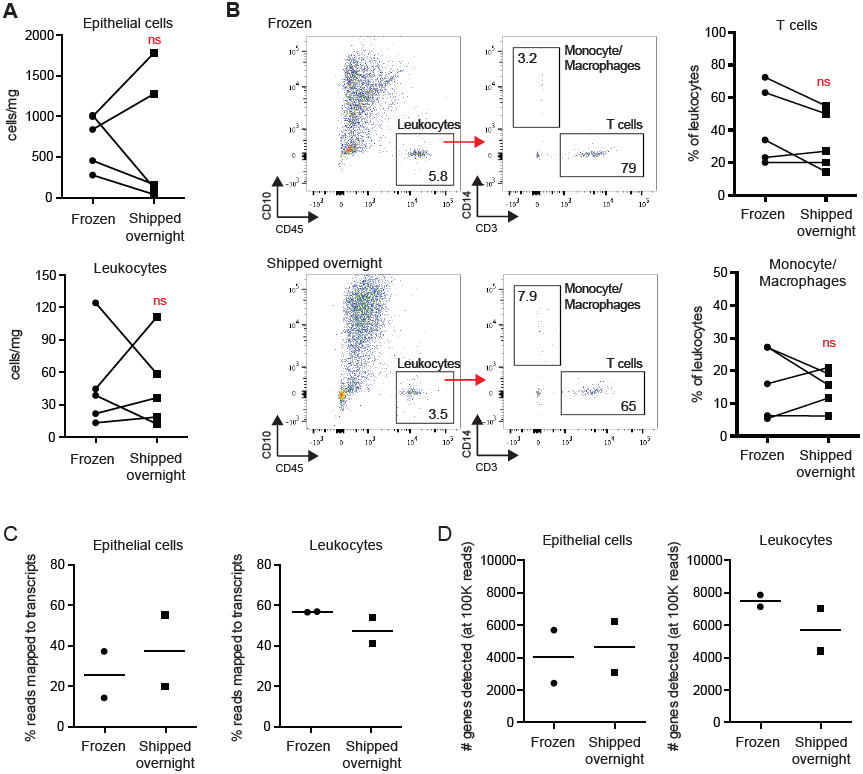
Overnight shipping of kidney tissues in storage solution or after cryopreservation leads to comparable cell yields. A) Cell yields by flow cytometry from tumor nephrectomy pieces either cryopreserved or shipped overnight on wet ice (n= 5 donors). Wilcoxon test. B) Flow cytometric detection of leukocyte populations within kidney tissue and quantification of percentage of T cells and monocyte/macrophages among leukocytes. Wilcoxon test. C,D) Percent of reads mapped to transcriptomes (C) and number of genes detected (D) in RNA sequencing data from of bulk leukocytes and epithelial cells from kidneys either cryo-preserved or shipped overnight.

Taken together, these results suggested the cryopreservation of intact tissue offers a feasible strategy for obtaining viable leukocytes and epithelial cells for batched flow cytometric and transcriptomic analyses.

### Dissociation of cryopreserved kidney tissue

Our studies so far showed that cryopreserved tissue yielded similar cell numbers, viability and RNA quality as samples shipped overnight in a storage solution, but had the advantage of a longer storage window, allowing the ability to batch samples for downstream analyses. Next we evaluated several variables in the dissociation method, as applied to cryopreserved kidney tissue.

Because it has been reported that fracture of endothelial cells in transplanted frozen tissue is dependent on thaw time (18), we compared rapid thawing of tissue in a 37°C water bath to slower thawing by allowing the tube to thaw on the benchtop (~6-7mins until thawed) and found no clear differences in cell yields (**Fig. 4a**, n=3).

**Figure 4.**
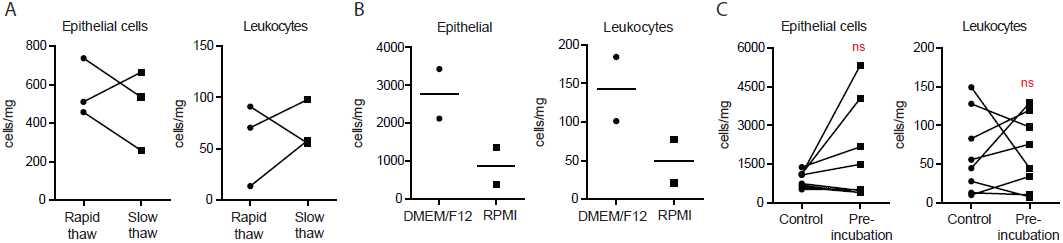
Enzymatic dissociation of cryopreserved kidney tissue. A) Flow cytometric quantification of cell yields from cryopreserved kidney tissue thawed rapidly or slowly and then dissociated (n=3 donors). B)) Epithelial cell and leuko-cyte yields after dissociation with liberase TL in DMEM/F12 or RPMI media (n=2 donors). C) Epithelial and leukocyte yields after dissociation with or without pre-incubation with liberase TL for 30 minutes on ice (n= 8 donors).

Tissue dissociation by mechanical disruption alone yielded few intact epithelial cells; therefore, we evaluated enzymatic digestion to obtain single cell suspensions. Experiments comparing tissue digestion using Liberase TL or collagenase P resulted in similar cell yields but demonstrated a transcriptomic signature of endotoxin exposure with collagenase P digestion (Table S2), consistent with prior observations(19,20). Liberase TL was therefore selected for enzymatic digestion. Digestion time courses ranging from 5-15 minutes indicated optimal cell yields with 12 minutes of enzymatic digestion with Liberase TL at 37°C (data not shown). Pilot experiments evaluating addition of the caspase inhibitor ZVAD or human serum albumin to the protocol did not show improvements over the current method (data not shown). Because nutrients in the digestion buffer may help preserve viability of the released cells, we compared digestion in RPMI and DMEM/F12. Digestion in DMEM/F12 demonstrated higher cell yields in two independent experiments; therefore digestion in DMEM/F12 was adopted for the protocol (**Figure 4b**, n=2).

Finally, we evaluated in several donors the utility of pre-incubating the tissue with Liberase TL at 4°C for 30 minutes to allow better tissue penetration but did not observe a consistent advantage over immediate digestion at 37°C (**Figure 4c**, n=8). The consensus protocol for tissue dissociation is described in the methods section.

### Application of method to LN kidney biopsies and urine samples

We evaluated 12 LN biopsy samples during the development of this protocol, including 4 LN biopsies processed by the final consensus protocol. LN biopsy sample yields ranged from ~50 leukocytes to over 3000 leukocytes per biopsy segment. We also evaluated urine samples from 6 LN patients, 4 of which yielded sufficient cells for analysis.

We implemented an 11-color flow cytometry panel developed to identify both epithelial cells and leukocyte subsets in kidney and urine samples. Flow cytometry of urine cells may be complicated by autofluorescence; however, we observed that the effect of autofluorescence varied between flow cytometric channels and could be minimized by optimized marker-fluorophore design. This panel clearly identified CD10+ epithelial cells and CD45+ leukocytes in both LN kidney biopsy samples and urine cells (**Figure 5**). Within the leukocytes, CD4^+^ and CD8^+^ T cells, CD14^+^ macrophages, and CD19^+^ B cells could be distinguished.

**Figure 5.**
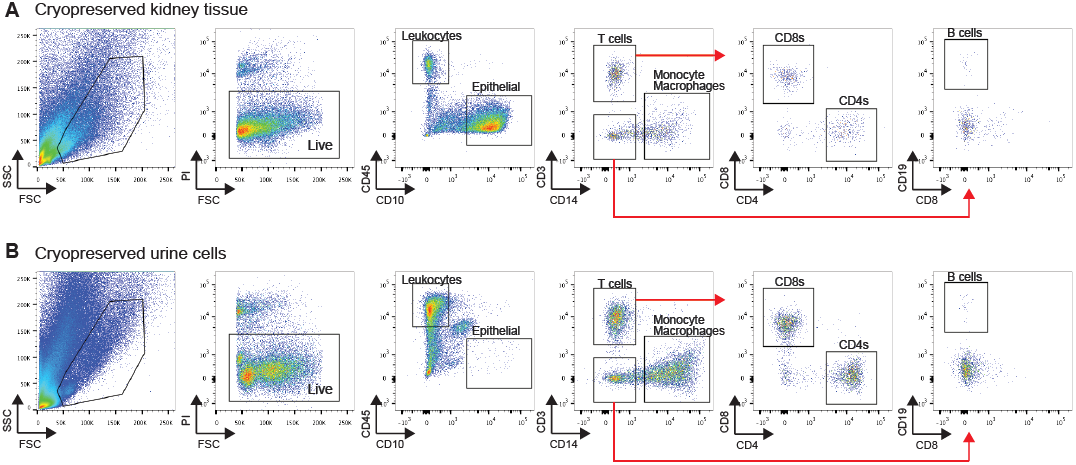
Flow cytometric analysis of lupus nephritis samples. A) Flow cytometry of cryopreserved lupus nephritis kidney biopsy tissue. B) Flow cytometry of cryopreserved lupus nephritis urine cells.

RNA-seq analysis of bulk-sorted cell populations from a LN biopsy revealed high quality library for CD45^+^ leukocytes and, to a lesser extent, for CD10^+^ epithelial cells, consistent with results obtained for tumor nephrectomy samples (**Figure 6a**). RNA-seq data were also successfully obtained from sorted bulk leukocyte populations from 3 LN urine cell samples, although the libraries were somewhat inferior to that observed for blood cells obtained from a different cohort of LN patients (**Figure 6b**). We compared the gene expression data from 3 sorted populations of urine cells – CD45^+^CD3^+^,CD45^+^CD3^-^ and CD45^-^ cells – to an external dataset, containing gene expression measurements from 360 different cell types (FANTOM5, http://fantom.gsc.riken.jp/5/)(21,22). We found that CD45^+^CD3^+^ cells most resembled different subsets of T cells; CD45^+^CD3^-^ cells were most similar to macrophages and monocytes; and in one out of two patients, gene expression data from CD45^-^ cells correlated best with renal cortical epithelial cells and renal proximal tubular epithelial cells (**Table S3**). These results indicate that transcriptomes derived from urine cells can indeed contain meaningful cell lineage information.

**Figure 6:**
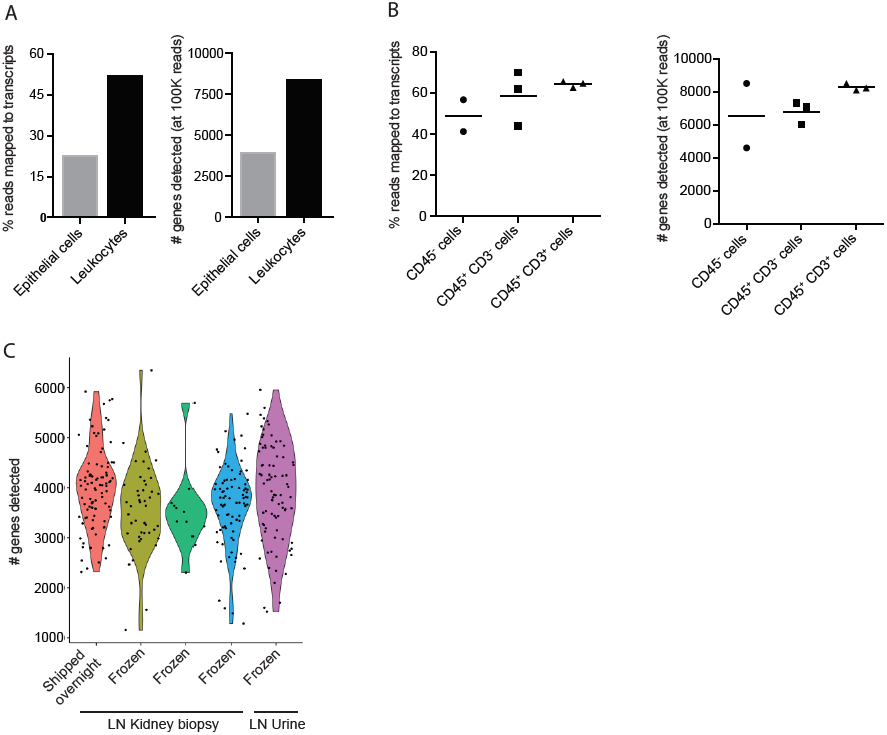
Transcriptomic data from LN samples. A) Quality metrics of bulk leukocyte and epithelial cell populations sorted from a cryopreserved LN kidney biopsy. B) Quality metrics of bulk cell populations (CD45^-^ cells, CD45^+^ CD3^-^ cells, CD45^+^ CD3^+^ cells) sorted from LN urine (n=3 donors, insufficient CD45-cells were obtained from 1 donor). C) Number of genes detected by single cell RNA sequencing of sorted individual CD45^+^ leukocytes from LN kidney biopsies from 4 different donors processed as indicated and from cryopreserved urine cells from a LN donor.

In recent years, RNA-seq data from single cells were shown to be informative in identifying known and novel subsets of immune cells, delineating pathways active under different conditions, and highlighting the differentiation course of cells *in vivo*. It is therefore of interest to determine if high quality single cell RNA-seq data can be obtained from LN kidney samples. To evaluate this, we sorted and profiled single CD45^+^ cells from 3 cryopreserved LN kidney biopsies and 1 LN biopsy shipped overnight on wet ice. The quality of the data was high in all 4 samples, with 3000-4000 genes detected per cell (**Figure 6c**). In addition, we profiled single CD45^+^ cells sorted from a cryopreserved urine sample from a LN patient; we found the quality of the resulting data similar to that of cells obtained from LN kidney samples (**Figure 6c**).

## DISCUSSION

A primary goal of the AMP RA/SLE network is to generate robust single cell transcriptomic data from kidney and urine samples obtained from a large cohort of LN patients. Assembly of such a cohort in a reasonable timeframe requires sample acquisition in a distributed research network across multiple sites. Processing of samples at one centralized analysis site offers the potential to reduce technical variability between samples and generate more robust analyses. The protocol described here offers a simple, efficient method to cryopreserve kidney tissue and urine samples immediately after acquisition for downstream analyses at a central processing site. This method has been approved by the AMP RA/SLE network and has been recently implemented to analyze 26 cryopreserved LN kidney biopsies in Phase 1.

We demonstrate that kidney tissue can be cryopreserved in a DMSO-containing solution, similar to what has been performed routinely for many years with peripheral blood mononuclear cells. Leukocytes from cryopreserved kidney show a phenotype similar to that seen in kidney samples shipped overnight on wet ice, and leukocyte yields are comparable between the two methods. Our results also demonstrate that cryopreservation of the intact tissue, rather than dissociated cells, substantially improves epithelial cell yields. We hypothesize that preserving epithelial cell attachment to basement membrane or adjacent cells during the cryopreservation process promotes retention of viable epithelial cells.

The ability to freeze tissue removes the burden of tissue dissociation from each of the acquisition sites, and allows for centralized tissue dissociation. Cryopreservation offers substantial logistical advantages over shipping tissues overnight on wet ice. Samples can be acquired and frozen on any day of the week without concern for availability of the processing site. In addition, the processing site does not need to be ready to immediately process a sample immediately upon arrival. Rather, cryopreserved samples can be shipped when convenient and processed centrally with flexible scheduling, allowing for design of well-matched batches of samples. Our RNA-seq results suggest a minor stress response following cryopreservation; however, the given the limited sample sizes here, additional studies are required to define characteristic expression changes with freezing, which may then be adjusted for computationally.

The dissociation protocol described here using enzymatic digestion with liberase TL combined with gentle mechanical disruption was evaluated with multiple readouts relevant to the pipeline, including assessments of cell yield, flow cytometric phenotype, and transcriptomics. We utilized transcriptomics to assess critical steps in the protocol to ensure that each variable selected did not negatively affect downstream transcriptomic quality. The current protocol allows for targeting of specified cell types, with an emphasis here on tissue-infiltrating leukocytes. Leukocytes can be specifically sorted out of the total cell suspension for bulk transcriptomic analyses or single cell transcriptomics. We demonstrate here high quality RNA-seq transcriptomes from both kidney and urine leukocytes. Current efforts are underway to further improve viability and transcriptome integrity of isolated kidney epithelial cells.

In summary, the method described here provides a robust method for isolation and transcriptomic analyses of single cells isolated from cryopreserved LN kidney and urine samples, which can be easily stored and transported. The limited demands for the staff at the acquisition sites allows for rapid dissemination to multiple sites and facilitates participation from sites with minimal technical laboratory support. We propose that this pipeline, developed for LN kidney tissues, may serve as a model for robust, centralized analyses of viable cells from kidney tissue acquired in multi-site clinical studies. More broadly, this strategy enables accumulation of a valuable biobank of tissues containing viable cells that can be used for multiple downstream analyses.

## ACKNOWLEDGEMENTS

We thank Brad Godfrey, J. Michelle Kahlenberg, Heather Ascani, and Michael Gurish for technical advice. We thank participating Lupus Nephritis Trials Network clinical sites and participants. This work was supported by the Accelerating Medicines Partnership (AMP) in Rheumatoid Arthritis and Lupus Network. AMP is a public-private partnership created to develop new ways of identifying and validating promising biological targets for diagnostics and drug development (http://fnih.org/what-we-do/current-research-programs/amp-ra-sle). Funding was provided by grants from the NIH (UH2-AR067676, UH2-AR067677, UH2-AR067679, UH2-AR067681, UH2-AR067685, UH2-AR067688, UH2-AR067689, UH2-AR067690, UH2-AR067691, UH2-AR067694, and UM2-AR067678).

---

**Figure S1.**
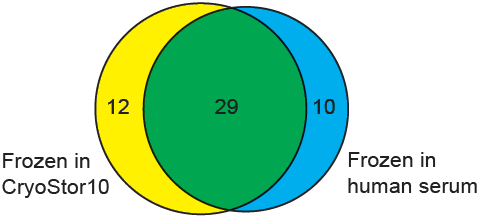
Overlap of genes upregulated (FDR:s 0.05) in leukocytes frozen in either CryoStor10 or human serum/10% DMSO, compared to fresh cells (2 samples of each condition were analyzed).

**Table S1.**
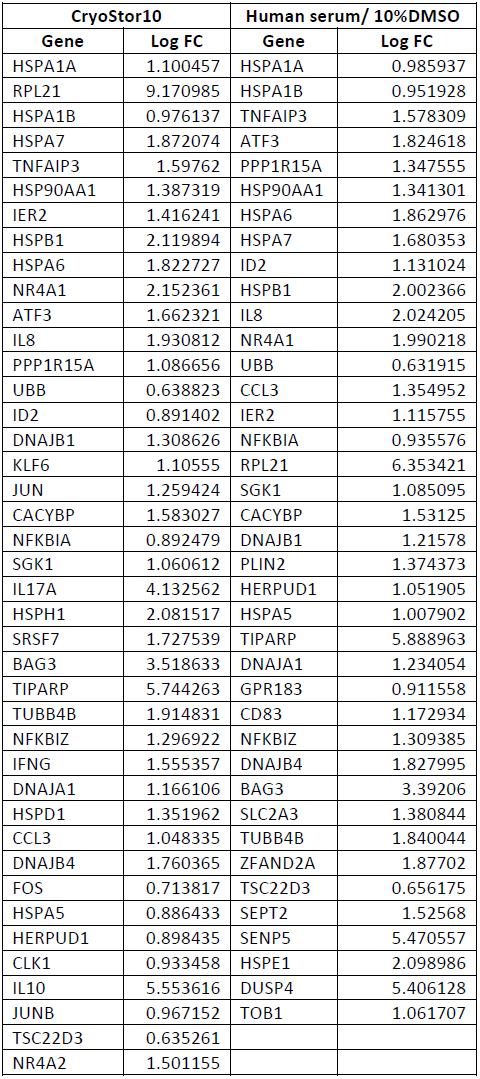
Genes found to be upregulated (FDR ≤ 0.05) in leukocytes, comparing frozen cells to fresh cells (2 samples of each condition were analyzed), considering two different freezing media.

**Table S2.** Genes upregulated (FDR ≤ 0.05) in leukocytes in tissue dissociated with collagenase P, compared to leukocytes obtained from tissue dissociated with Liberase TL. * indicates genes previously reported to be induced by LPS(19).

| Gene | Log FC |
| --- | --- |
| IL1B* | 1.714549341 |
| IL8* | 2.124210362 |
| NFKBIA* | 1.096730002 |
| NFKBIZ* | 1.553312222 |
| CCL3* | 1.29642886 |
| TYROBP | 0.88167085 |
| NKG7 | 0.65205601 |
| METTL21A | 2.84074085 |
| CCL4L1* | 1.021842283 |
| CCL4L2 | 1.021852219 |
| IER3 | 1.941868195 |
| CTSS | 0.562055522 |
| ZFP36* | 0.682632534 |

**Table S3.**
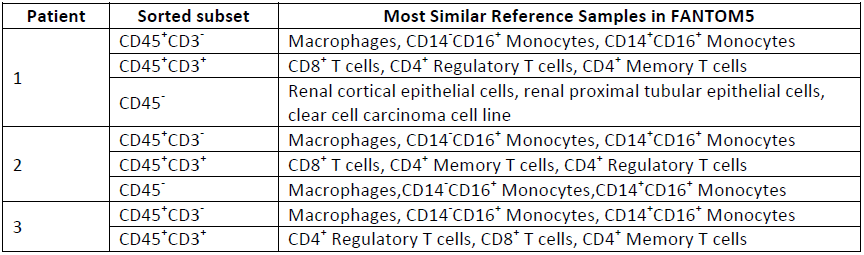
The results of comparing sorted urine cell populations from 3 LN patients to an external data set (FANTOM5). For each sorted cell population, the table lists the top 3 FANTOM5 samples with which it had the highest correlation score.

